# JIP2 haploinsufficiency contributes to neurodevelopmental abnormalities in human pluripotent stem cell-derived neural progenitors and cortical neurons

**DOI:** 10.1101/196535

**Authors:** Reinhard Roessler, Johanna Goldmann, Chikdu Shivalila, Rudolf Jaenisch

## Abstract

Phelan-McDermid syndrome (also known as 22q13 deletion syndrome) is a syndromic form of autism spectrum disorder and currently thought to be caused by heterozygous loss of SHANK3. However, patients most frequently present with large chromosomal deletions affecting several additional genes. We used human pluripotent stem cell technology and genome editing to further dissect molecular and cellular mechanisms. We found that loss of JIP2 (MAPK8IP2) may contribute to a distinct neurodevelopmental phenotype in neural progenitors (NPCs) affecting neuronal maturation. This is most likely due to simultaneous misregulation of JNK proteins, leading to impaired generation of mature neurons. Furthermore, semaphorin signaling is compromised in patient NPCs and neurons. Pharmacological stimulation of neuropilin receptor 1 (NRP1) rescued impaired semaphorin pathway activity and JNK expression in patient neurons. Our results suggest a novel disease-specific mechanism involving the JIP/JNK complex and identify NRP1 as potential therapeutic target.

## Introduction

22q13 Deletion Syndrome (22qDS) is commonly caused by deletions in the q-terminal end of chromosome 22. Characteristic symptoms of 22qDS include developmental delay, intellectual disability and severe delay or complete absence of speech (Phelan, 2008). 22qDS is currently seen as synaptopathy, predominantly caused by heterozygous deletion of the gene *SHANK3*. The importance of SHANK3 protein for functional synapses has been robustly established (Durand et al., 2007; Jiang and Ehlers, 2013; Mei et al., 2016; Peça et al., 2011; Uchino and Waga, 2013). Very recently SHANK3 has been functionally implicated in subcellular signaling pathways, such as the mTOR-pathway, and it has been shown that SHANK3 depletion impairs intracellular signaling (Bidinosti et al., 2016; Shcheglovitov et al., 2013).

However, the contribution of additional, concomitantly deleted genes in this disease is largely unknown. The vast majority of patients nonetheless present with highly variable chromosomal deletions that can effect several genes located up or downstream of *SHANK3* (Bonaglia et al., 2011; Sarasua et al., 2014). One interesting candidate gene commonly co-deleted in 22qDS is *MAPK8IP2* (aka *JIP2*). JIP2 is know to be a crucial scaffolding protein that facilitates the activities of MAP kinase pathway proteins including MEKs, ERKs and JNKs (Whitmarsh, 2006). Regulated activation of the MAPK pathway, and in particular correct regulation of JNK proteins, has been shown to be crucial for neuronal differentiation as well as for neuron function (Coffey, 2014a; Tiwari et al., 2011). Interestingly, Tiwari et al. (2011) showed that JNK inhibition results in impaired neuronal differentiation of mouse embryonic stem cells (ESCs). It is therefore conceivable that JIP2 haploinsufficiency also impacts early human neurodevelopment. Supporting this notion, imaging studies of a small group of Phelan-McDermid patients revealed a neurodevelopmental phenotype predominantly effecting the formation of the cerebellum (Aldinger et al., 2013). Crucially, such a morphological phenotype has not been observed in *Shank3* knockout mice.

Given the complexity of the deletions, we hypothesize that, while a small number of patients lacking only *SHANK3* have been diagnosed, 22qDS may frequently not only be the consequence of heterozygous loss of *SHANK3* but might additionally be linked to loss of *JIP2*. In this study we present evidence that both SHANK3 and JIP2 contribute to disease-specific phenotypes that occur at distinct neurodevelopmental stages. Based on these observations we propose a disease mechanism that involves the MAPK pathway and the regulatory function of JNK proteins as well as impaired mTOR pathway activity. We believe that the realization of this complex neurodevelopmental and functional phenotype is crucial for successful development of potential pharmacological interventions.

## Results

### Patient-specific iPSCs show impaired neuronal maturation

Patient-specific iPSCs and ESCs were differentiated into NPCs and mature cortical neurons and immunoflourescence and immunoblot analysis was used to characterize basal identities (Fig.1A-C). Morphological and molecular profiling revealed that both patient and control PSCs could efficiently generate cortical neurons. Both cell populations sequentially differentiated into cells expressing NPC markers such as PAX6 and NESTIN and continued *in vitro* maturation resulted in neurons expressing pan-neuronal markers such as TUJ1 and MAP2. Comparing patient and control neurons, we observed the expected reduction of protein levels for SHANK3 and JIP2 as well as a severe reduction of neuronal markers such as DCX and NeuN (Fig.1C&D). JNK proteins were only weakly induced in patient neurons as compared to control neurons. Gradual induction of JNK during differentiation of mouse ESCs into neurons has been documented during neural induction (Tiwari et al., 2011). Strikingly, impaired up-regulation of JIP2 in patient neurons coincided with failed induction of JNK proteins (Fig.1C&D). This observation prompted us to investigate electrophysiological properties of mutant cells as an additional criterion for neural maturation. We performed multi-electrode array (MEA) analysis (Maestro, Axion) comparing control neurons and 22qDS neurons during *in vitro* maturation for up to 95 days. Comparative quantification of the neuron content in both conditions did not show any significant deviation (Fig.1E). To ensure equal cell density of neurons plated we counted the total number of cells (approx. 120.000 cells/well) and evaluated the neuron content by immunocytochemistry of parallel (Fig.1F-G) cultures and Western Blot analysis (data not shown). Average spike quantification over a 3-month period revealed a significant impairment of *in vitro* maturation in patient-derived neurons (Fig.1I). While control neurons gradually increased their spontaneous activity only small surges were observed in patient neurons. Comparative raster plots across 64 individual electrodes at day 71 of differentiation clearly showed higher activity for control neurons (Fig.1J). Quantification of active electrodes across differentiation experiments comparing control and patient neurons also revealed reduced over-all activity in 22q13 neurons at later stages (Fig.1K). By analyzing the mean spike rate during *in vitro* maturation we found a gradual increase over time for control neurons (Fig.1L). Together with increasing numbers of active electrodes this observation indicates an increase in network activity in the control cultures. During the differentiation time-course spike rates remained low for patient neurons with significant reduction particularly in late stage mutant neurons. Thus, our results indicate severely impaired network activity in patient neurons.

**Figure 1:**
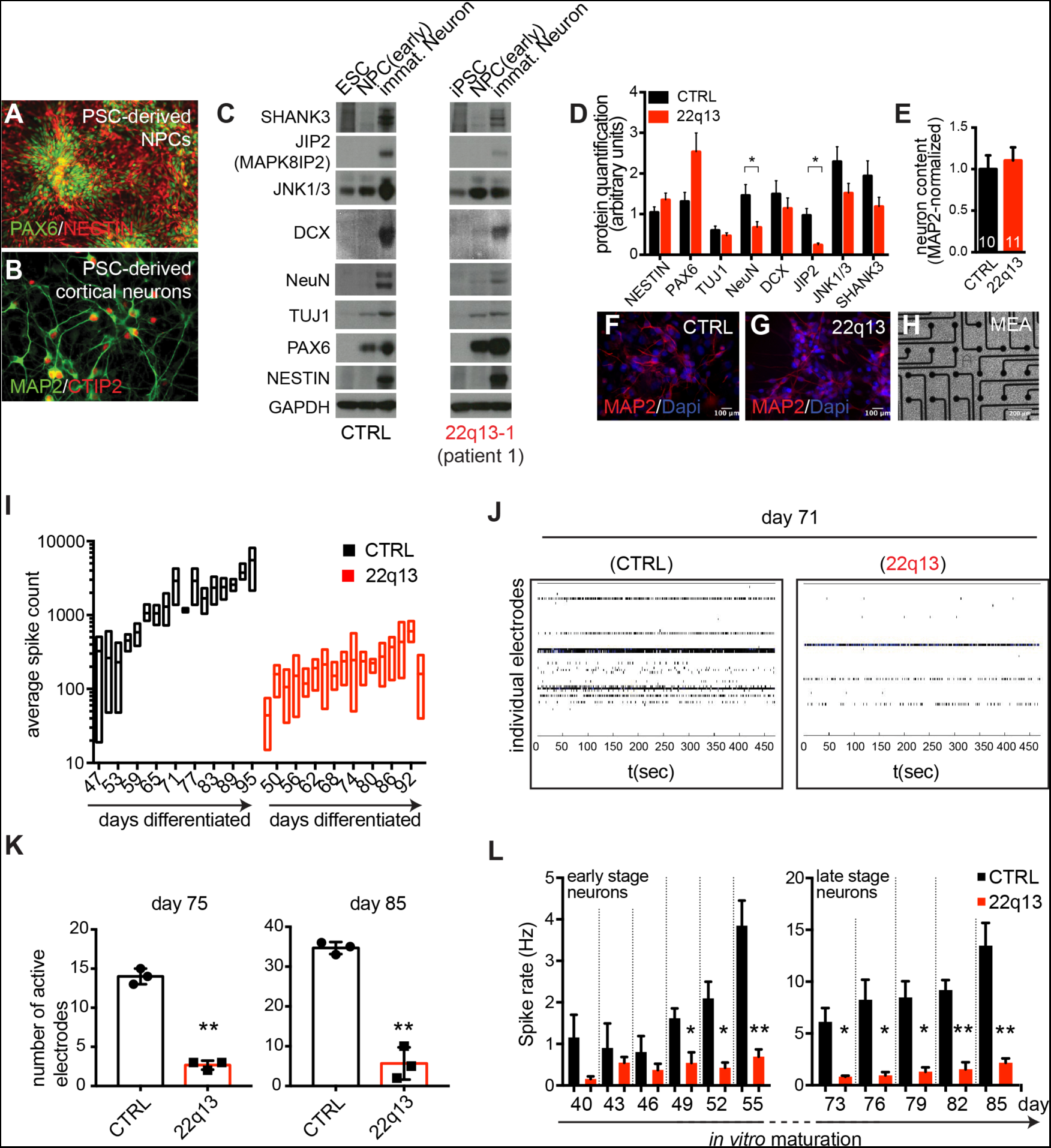
Patient-specific iPSCs show impaired neuronal maturation. (A) Immuno-staining for PAX6 and NESTIN in PSC-derived NPCs. (B) Immuno-staining of PSC-derived cortical neurons for MAP2 and CTIP2. (C) Comparative protein analysis for SHANK3 and JIP2 as well as for JNK proteins, NPC- and pan-neuronal markers for CTRL and patient lines. (D) Quantification of protein levels in neurons. Data are shown as mean ± SEM (n=3). Two-tailed unpaired t-test, h^*^ p < 0.05. (E) Quantification of normalized neuron content plated on multi-electrode arrays. Data are shon as mean±SEM (n=10/11 images per condition, respectively). (F&G) MAP2 immuno-staining of age/culture-matched (not density-matched) neurons used for MEAs. (H) Representative phase-contrast image showing neurons seeded on 64 electrodes/well. (I) Quantification of average spike count during *in vitro* maturation. Average of 3 wells (3 independent cultures) for 5min measurement intervals are shown as box plots ± SEM. (J) Comparative raster plots as recorded on day 71 of differentiation. Every line in individual plot represents one electrode. Every vertical dash represents a detected field potential. Recording interval: up to 450 sec. (K) Quantification of number of active electrodes at day 75 and day 85. Data are represented as mean ± SEM. Two-tailed unpaired t-test, ^**^ P < P< 0.005. (L) Quantification of average spike rate in Hz at early and late stage differentiation. Data are represented as mean ± SEM. Two-tailed unpaired t-test, ^*^ p < 0.05, ^**^ p < 0.005

Postsynaptic density proteins establish tightly packed aggregates to facilitate efficient synaptic transmission. Furthermore, these structures are directly linked to intracellular downstream signaling pathways (Suppl. Fig.1A).Protein analysis in 3 month old neurons confirmed a severe reduction of SHANK3 in the 22qDS mutant cells associated with reduced levels of direct and indirect binding partners of SHANK3 while pan-neuronal markers appear to be unchanged (Suppl. Fig.1B). In addition, we observed reduced protein levels of AKT and mTOR and their phospho-variants in patient neurons. The AKT/mTOR pathway has recently been associated with Phelan-McDermid Syndrome (Bidinosti et al., 2016). We also detected reduced levels of Homer and PI3K enhancer (PIKE), which mediate signal transduction to PI3-Kinase and eventually trigger the activation of the mTOR complex (Suppl. Fig.1C). This result confirms that loss of SHANK3 not only leads to impaired synaptic transmission but also influences central intracellular pathways in mature neurons.

### Mutant iPSC-derived neural progenitors reveal a neurodevelopmental phenotype

Our initial observation that 22qDS neurons show severely impaired *in vitro* maturation led us to explore whether neurodevelopmental defects might be detectable already earlier during the differentiation process and consequently result in functional differences. For this we genetically engineered fluorescent NPC reporter cell lines for the patient and control background that would allow purification of a defined cell population. We used CRISPR/Cas9 gene targeting to integrate a GFP reporter cassette into the endogenous locus of *PAX6*, a well characterized NPC marker (Georgala et al., 2011; Osumi et al., 2008) (see Fig.2A for targeting strategy). Correctly targeted clones were identified by puromycin selection and Southern Blot assays (Fig.2B) and differentiation of these clones into NPCs activated the PAX6-GFP reporter in cells with typical NPC morphology (see representative immuno-staining in Fig.2C). Clones with CRISPR/Cas9 induced double strand breaks in the second allele (PAX6^GFP/-^) resulted in largely GFP negative cell populations upon differentiation and were therefore excluded from further analyses (suppl. Fig.2A&B). Separation of GFP positive and GFP negative populations by FACS (Fig.2D and suppl. Fig.2B) enabled the analysis of global gene expression in a homogenous NPC population. Expression profiling identified a large set of differentially expressed genes comparing the mixed population with the GFP positive and GFP negative fraction (Fig.2E and suppl. Fig.2C). More specifically, NPC markers were up regulated in the GFP positive fraction (Fig.2F), while genes up-regulated in the GFP negative fraction were identified as neural crest markers (data not shown). Gene ontology analysis revealed the up-regulated population of genes in the NPC fraction as genes characteristic for forebrain progenitors (strongly enriched GO terms are e.g. telencephalon and forebrain development, axon guidance, neuron differentiation, etc.) (Fig.2G). This result confirms PAX6 as robust NPC marker that can be used to purify a defined progenitor population. More detailed transcript analysis showed that NPC markers such as NESTIN, MSI1, ASCL1 and several SOX family members were equally expressed in purified control and patient NPCs. (suppl. Fig.2D&E).

**Figure 2:**
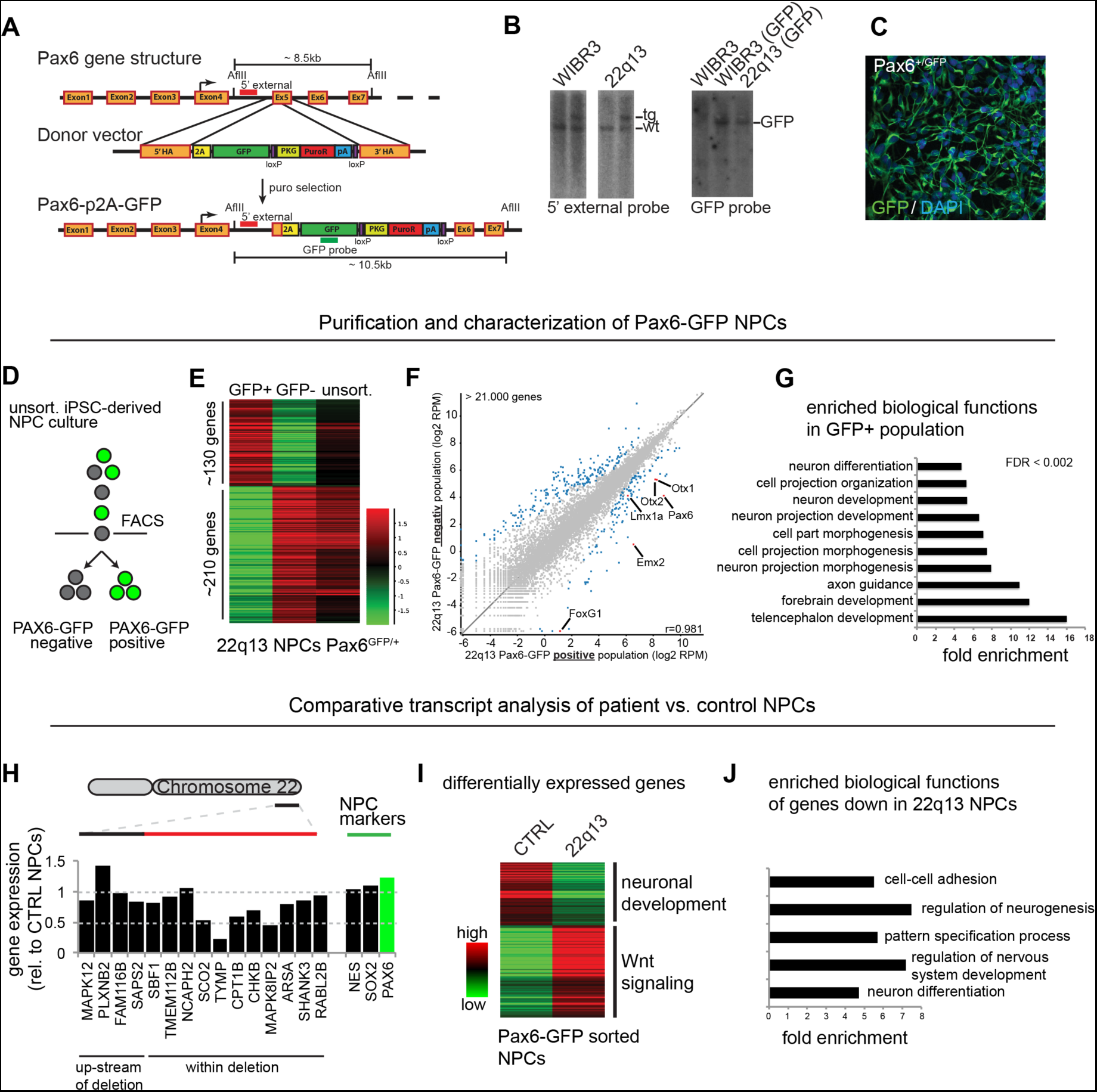
Patient neural progenitors show a neurodevelopmental phenotype. (A) CRISPR/Cas9 targeting scheme for fluorescent PAX6-GFP reporter. (B) Southern Blot analysis for targeted genomic locus showing 5’ external and internal probe for control and patient specific iPSC lines. (C) Representative immuno-staining for GFP upon NPC differentiation. (D) FACS scheme separating three distinct cell populations. (E) Heat map of differentially expressed genes comparing GFP positive, GFP negative and unsorted cell populations. (F) Scatter plot representing global gene expression levels. Blue dots represent up-regulated genes in GFP+ and GFP-fraction, respectively. Red dots highlight selected NPC marker genes in the GFP+ fraction. (G) GO term analysis of genes up-regulated in GFP+ fraction. (H) Quantitation of relative gene expression (rel. to CTRL NPCs) across patient-specific deletion. Selected NPC marker gene expression is shown as reference for PAX6-GFP sorted populations. (I) Differentially expressed genes comparing GFP+ cells form controls and patient line. (J) GO term analysis of genes down-regulated in patient NPCs (GFP+ cells).

Having identified specific NPC signatures in control and patient Pax6-GFP sorted cells we used this system to compare the expression profiles between purified control and patient NPCs to identify potentially disease relevant differences. RNA-Seq analysis showed that relative expression levels of genes within the deletion were largely reduced in patient NPCs and fluctuated around a 0.5 fold expression while expression of a set of NPC markers showed a high level of similarity (Fig.2H and suppl. Fig.2F). Global gene expression analysis identified a large set of genes (approx. 300) that were differentially expressed in purified patient NPCs (Fig.2I) and gene ontology analysis classified the down-regulated fraction as genes with biological functions such as ‘regulation of neurogenesis’, ‘regulation of nervous system development’ and ‘neuron differentiation’ (Fig.2 J). This result provides evidence that phenotypic changes in 22qDS NPC already occur before mature synapses have been established and underlines the notion that mutations in SHANK3 may not be the sole factor determining the pathology in the majority of Phelan-McDermid syndrome patients.

### Isogenic ESC-derived neural cell types recapitulate patient-specific phenotypes

Comparing different iPSC lines in order to evaluate disease relevant phenotypes can be particularly challenging as heterogeneous genetic backgrounds may drastically mask subtle cellular phenotypes. Recent advances in genome editing, however, have enabled the generation of isogenic mutant lines that are genetically identical except for the disease relevant deletion or mutation (Li et al., 2013; Soldner et al., 2011). To generate isogenic ESC lines that carry a similar deletion as the patient-specific iPSC line used above we expressed Cas9 protein and deletion-flanking gRNAs in the WIBR#3 hESC line (see Fig.3A for targeting scheme). We obtained clones that carried a heterozygous deletion of approximately 93kb of the long arm of chromosome 22. PCR analysis across or flanking the deletion site identified three heterozygous clones that recapitulated a patient-specific deletion (Fig.3B). Sequence analysis of the “wild type” allele showed that the 5’ targeting site contained only a single base pair deletion upstream of the *JIP2* gene (Fig.3C). This frame shift mutation however did not impact heterozygous expression of JIP2 (Fig.3F&G). Sequencing of the 3’ targeting site on the other hand indicated several INDELs in the isogenic deletion lines (Fig.3C). We found deletion-specific reduction of SHANK3 particularly affecting individual isoforms in isogenic Δ22 lines (Fig.3F&G). Differentiation into NPCs appeared to be normal as assessed by immunostaining for NESTIN and PAX6 (Fig.3D&E). Subsequent differentiation into cortical neurons resulted in equal expression of the pan-neuronal marker TUJ1 in all compared lines (WIBR#3, WIBR#3_Δ22-1, WIBR#3_Δ22-2 and 22q13 iPSCs). Expression of JIP2 and the JIP binding partner JNK 1/3 was, however, reduced in all deletion lines (Fig.3 F&G). This result shows that genetically engineered 22qDS lines reliably recapitulate molecular phenotypes as observed in patient-specific cell types. In addition we observed reduced levels of DCX in isogenic ESC-derived neurons (Fig.3F&G)), confirming the neurodevelopmental molecular phenotype described earlier in iPSC-derived neurons.

**Figure 3:**
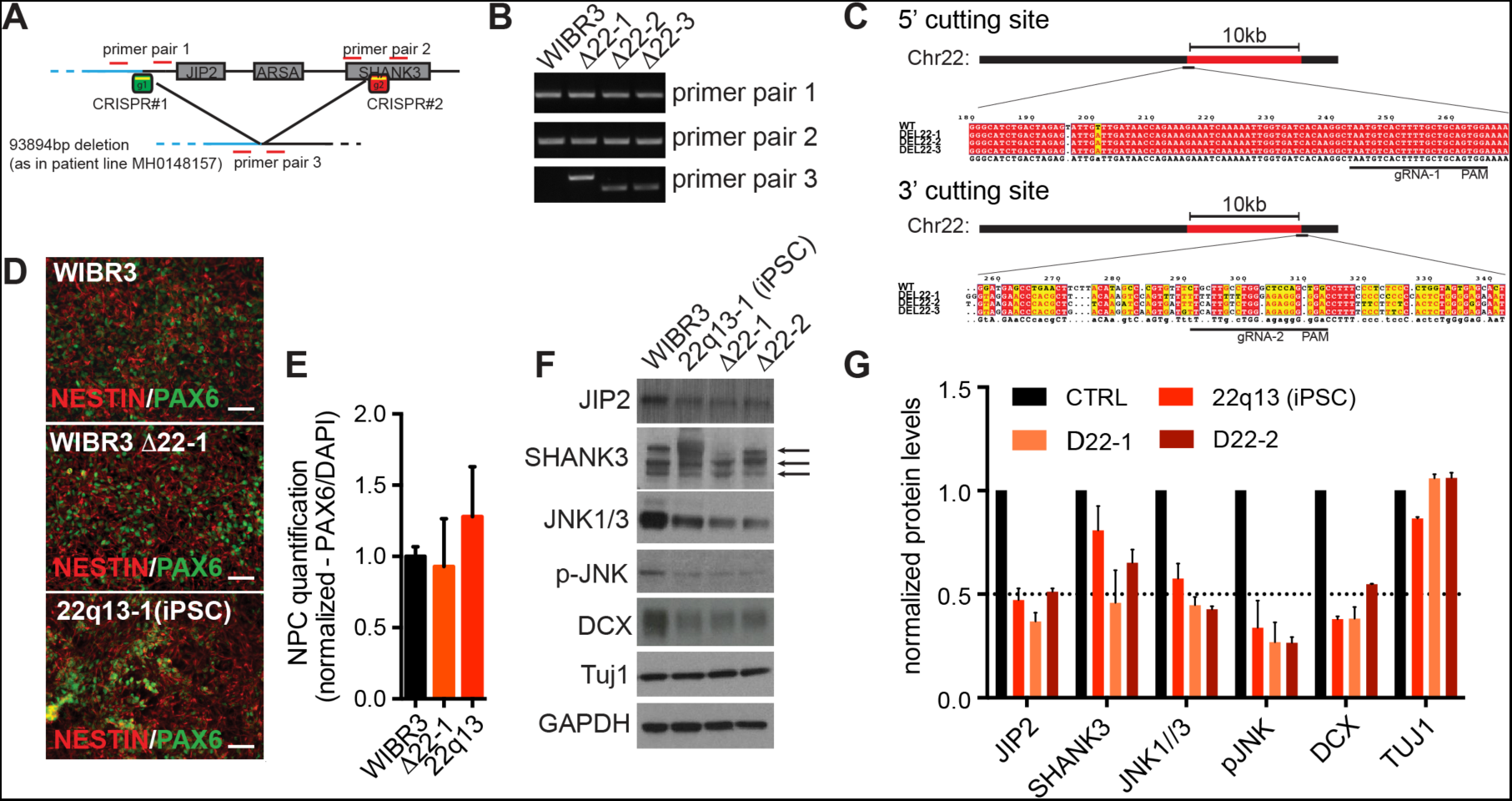
Isogenic ESC-derived neural cell types recapitulate patient-specific phenotypes. (A) CRISPR/Cas9 targeting scheme to introduce patient-specific deletion in WIBR#3 ESC line. (B) PCR assay shows untargeted control line and three heterozygous deletion lines. (C) Genomic DNA sequence of CRISPR target sites flanking the deletion. (D) Immuno-staining for control, isogenic deletion and iPSC lines after NPC differentiation. (E) Quantification of NPC fraction. Normalized data are shown as mean±SEM (n=4). (F) Comparative protein analysis of control-, isogenic deletion- and iPSC-lines upon neuronal differentiation. Arrows indicate three isoforms of SHANK3. (G) Quantification of immunoblots shown in (F). Data are represented as mean±SEM (n=3).

### Pharmacological activation of the neuropilin 1 receptor recues JNK expression in maturing patient neurons

Molecular and electrophysiological phenotypes have been described before in the 22qDS mutant cells. However, the contribution of *JIP2* haploinsufficiency to 22qDS has not been investigated in human neurons. Maturation of cortical pyramidal neurons depends on robustly established semaphorin signaling which subsequently activates JNK proteins (Calderon de Anda et al., 2012). Moreover it has been established that JIP and JNK proteins form functional complexes in order to facilitate kinase activities (Koushika, 2008; Kukekov et al., 2006; Mooney and Whitmarsh, 2004). We therefore hypothesized that JIP2 haploinsufficiency and consequently reduced JNK activity might contribute to impaired neuronal maturation. In order to elucidate potentially underlying molecular pathways we characterized known signaling mediators within the semaphorine pathway. Semaphorins are secreted guidance cues that bind to neuropilin receptors such as NRP1 (Maden et al., 2012; Pasterkamp, 2012; Tran et al., 2009). NRP1 activates the downstream kinase TAOK2 (thousand-and-one amino acid 2 kinase) which in turn phosphorylates and activates JNK proteins and tightly regulated JNK activity is know to be crucial for e.g. neuronal maturation and axon guidance processes (see Fig.4A for semaphorin pathway scheme) (Coffey, 2014b; Koushika, 2008). RNA-Seq analysis revealed that, while JIP2 is reduced in patient-derived NPCs, expression levels of TOAK2 and JNK1 are similar when comparing control and patient NPC transcript levels. Strikingly, we observed an increase in NRP1 transcript in 22qDS NPCs (Fig.4B). Except for DCX only minor transcript reductions were found in a set of direct JNK target genes (Fig.4B). However, protein analysis in neurons showed a decrease of JIP2, TOAK2, JNK1/3 and cJun (Fig.4C). These observations are consistent with the possibility that the function of JNK proteins rely on JIP binding particularly in differentiating neurons. In support of this notion we find that pharmacological stimulation of the NRP1 receptor by recombinant SEMA3A a) reverses up-regulation of NRP1 transcripts in patient neurons and restores control levels after 2h of *in vitro* exposure and b) rescues the expression levels of TOAK2 and JNK but did not stimulate JIP2 expression (Fig.4D).

**Figure 4:**
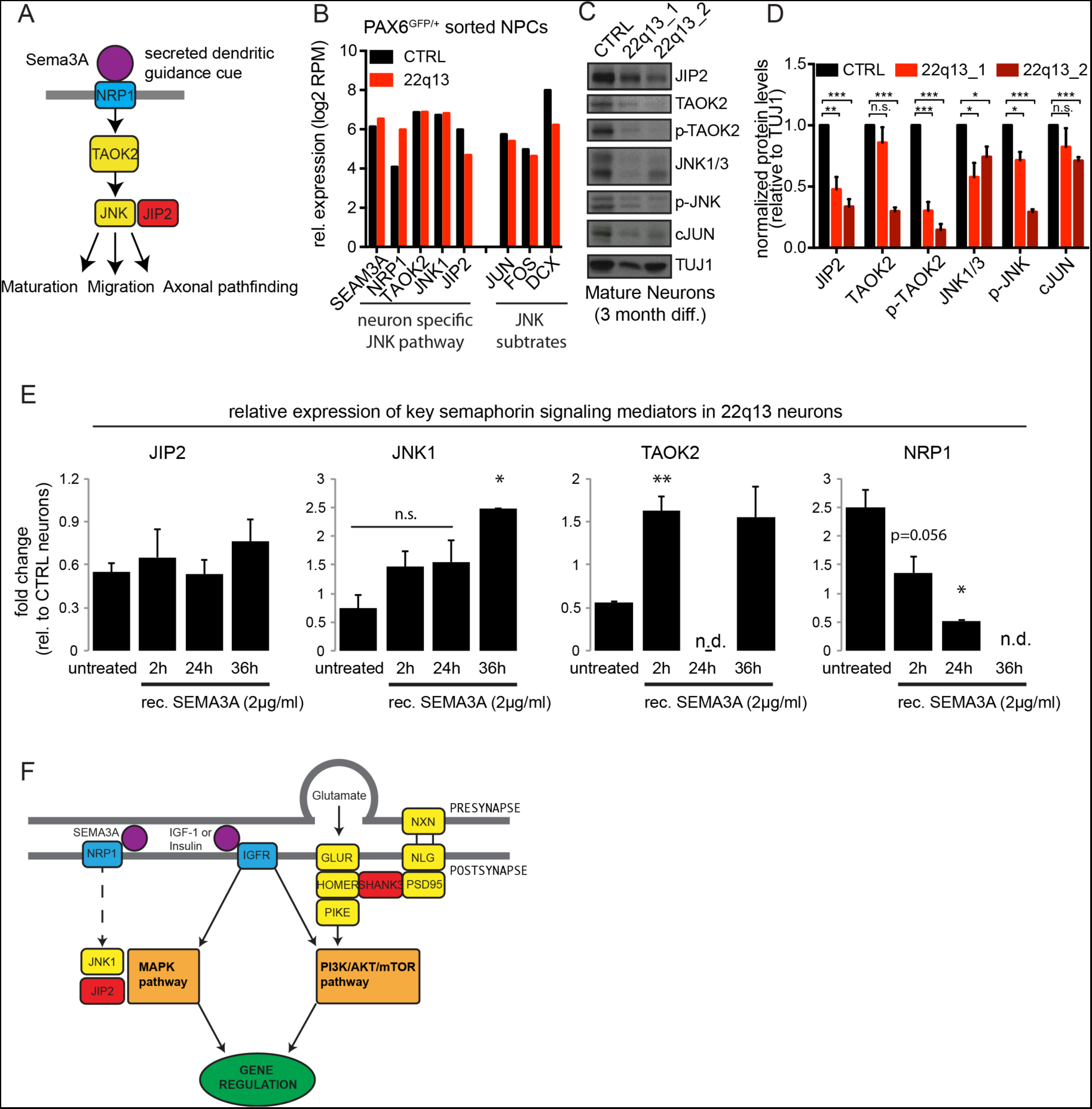
Pharmacological activation of the neuropilin 1 receptor recues JNK expression in maturing patient neurons. (A) Mechanistic scheme of semaphorin pathway in neurons. (B) Comparative transcript analysis (RNA-Seq) of key factors of semaphorin pathway and JNK target genes. (C) Protein analysis of untreated mature neurons derived from control and two independent patient iPSC lines. (D) Quantification of immunoblots shown in (C). Data are represented as mean±SEM (n=3). (E) Transcript analysis (qRT-PCR) of key factors of the semaphorin pathway upon treatment with rec. SEMA3A. Data are represented as mean±SEM (two biological replicates including three technical replicates each). Two-tailed paired t-test, ^*^ p < 0.05, ^**^ p < 0.005. (F) Mechanistic molecular model for 22q13 deletion syndrome.

## Discussion

We used pluripotent stem cells to study molecular and cellular phenotypes in NPCs and cortical neurons of 22qDS patients. Detailed analyses of defined neurodevelopmental stages suggested a novel disease-relevant mechanism that manifests in NPCs and negatively impacts neuronal maturation. Loss of JIP2 may impact expression levels of a large set of neurodevelopmental genes. Mechanistically this can be linked to impaired JNK complex function, and pharmacological stimulation of the semaphorin pathway rescues JNK expression. Our study adds mechanistic insight to this complex neurodevelopmental disease beyond SHANK3 haploinsufficiency.

22qDS has recently been connected to impaired synaptic transmission and disrupted intracellular signaling (Bidinosti et al., 2016; Lu et al., 2016; Shcheglovitov et al., 2013; Yi et al., 2016). Both cellular phenotypes place SHANK3 at the center of the underlying molecular mechanisms. While these findings significantly improve our understanding of this complex disorder it is crucial to note that the vast majority of patients have large chromosomal deletions, which encompass many adjacent genes consistent with mechanistic causes likely being more complex than currently recognized.

Human pluripotent stem cells provide a powerful tool to study such complex disorders, particularly if pathogenic genetic mutations have been identified (Soldner and Jaenisch, 2012; Soldner et al., 2016; Takahashi and Yamanaka, 2013). Here we combined human PSC technology and genome editing techniques to further study and dissect disease-relevant mechanisms. We confirmed findings that identified SHANK3 as a crucial contributor to symptoms associated to 22qDS. Loss of SHANK3 impaired intracellular signaling, in particular the AKT/mTOR pathway, and led to reduced synaptic activity in neurons *in vitro*. Recently, MEA recordings from Shank3 knockout mouse cortical neurons showed a reduction in network activity (Lu et al., 2016). Our findings in human cortical neurons confirm this observation. However, impairments seen in a patient neurons appear to be more pronounced. The more severe cellular phenotype seen in patient-specific neurons maybe be due to contributions of additional genes that are most commonly deleted in 22qDS.

We found that deletion of *JIP2* (aka *MAPK8IP2 or IB2*) likely contributes to an early neurodevelopmental phenotype that can already be detected in neural progenitors. While the role of JIP2 in ASD-type diseases is poorly understood, genetic mouse models support a relevant role of this MAPK pathway-related scaffold protein (Giza et al., 2010). Moreover, JIP proteins have been identified as direct binding partners of JNK proteins, which are know to be involved in neuronal maturation and proper neuron function (Coffey, 2014b; Koushika, 2008; Mooney and Whitmarsh, 2004; Whitmarsh, 2006). We found that loss of JIP2 coincided with reduced JNK expression in NPCs and mature neurons and detailed analysis of purified iPSC-derived NPCs identified a large set of neurodevelopmental genes to be misregulated. Previous studies described the pivotal role of JNK proteins during *in vitro* differentiation of mouse ESCs towards terminally differentiated neurons. JNK complexes specifically bind and activate the promoters of neurodevelopmental genes and inhibition of JNK leads to severely impaired neuronal differentiation, identifying JNK proteins as epigenetic regulators during neuronal maturation (Tiwari et al., 2011). We hypothesize that loss of JIP2 negatively impacts complex formation with JNK, which consequently contributes to a neurodevelopmental defect. Notably, neuropilin receptor 1 (NRP1), a semaphorin-specific receptor in developing neurons, was found to be up-regulated in patient NPCs and neurons. This specific increase in NRP1 might be a compensatory mechanism in order to maintain activity levels of the semaphorin pathway as needed for properly regulated neuronal maturation. Conversely, we also found that pharmacological stimulation of NRP1 restored JNK levels in neurons and NRP1 itself returned to control levels. This suggests a regulatory feedback loop in which JNK impacts NRP expression and vice versa. However, genetically controlled knockout experiments are needed to ascertain this theory. We did not observe significant up-regulation of JIP2 upon NRP1 stimulation. While it is possible that expression of the wild-type allele is increased, a more complex gene regulation may prevent specific induction of JIP2 in patient cells.

Fig.4F depicts a mechanistic model where loss of JIP2 impairs neuronal maturation by impacting the function of JNK and the MAPK pathway early in neurodevelopment. In addition, loss of SHANK3 results in impaired synaptic transmission and alleviated activity of the AKT/mTOR pathway later in mature neurons. This model also takes into consideration that IGF-1 (and insulin) triggers the activation of the AKT/mTOR pathway as well as the MAPK pathway and is therefore is able to ameliorate some 22qDS phenotypes. Earlier studies have shown that IGF-1 treatment improves cellular and behavioral phenotypes and even disease specific symptoms in patients (Bozdagi et al., 2013; Kolevzon et al., 2014; Shcheglovitov et al., 2013).

Modeling complex neurological diseases such as 22qDS is challenging and require human disease relevant systems as animal models might not be optimal to understand specific and potentially subtle phenotypes of human disease. Further investigation is needed to carefully dissect the independent roles of SHANK3 and JIP2. While previous studies focused on iPSC lines from patients that did not present with *JIP2* deletions it would be useful to engineer isogenic ESC lines that carry a deletion for either *SHANK3* or *JIP2*. Such a genetically defined approach would provide robust answers regarding the involvement of both candidate genes. Our study adds important insight in the understanding of underlying mechanistic processes in 22qDS and suggests neuropilin receptors as additional therapeutic targets.

## Experimental Procedures

### IPSC induction and neural differentiation

22qDS patient fibroblasts were acquired from the RUCDR cell repository (http://www.rucdr.org/1) and reprogrammed using the Stemgent RNA reprogramming kit (Warren et al., 2010). Briefly, patient fibroblasts were plated on NuFF feeder cells in varying densities to determine the most efficient reprogramming condition and transfected daily with the reprogramming cocktail for about two weeks (see Stemgent instructions). Clones were picked manually and expanded on inactivated mouse fibroblasts. WIBR3, a human ESC line described previously (Lengner et al., 2010), was used as control line. All cell lines used and generated are summarized in table 1. For neural induction and cortical neuron differentiation PSCs were exposed to dual SMAD inhibition combined with established optimized protocols for cortical in vitro differentiation (Chambers et al., 2009; Shi et al., 2012a, 2012b). PAX6-GFP reporter lines were differentiated for up to 4 weeks and were subjected to FACS upon up-regulation of GFP. Mature cortical neurons were derived from PSC-derived NPC by extended differentiation in N2/B27 medium for up to 3 month. Differentiation stages were identified by morphological characteristics and cell type specific marker expression.

### Electrophysiological characterization of PSC-derived neurons

To analyze electrophysiological properties of maturing neurons we used Axion Multi-Electrode-Arrays (https://www.axionbiosystems.com/). Electrode-containing wells were coated with poly(ethyleneimine) solution (Sigma-Aldrich, P3143) and Matrigel (VWR, #47743). Approx. 120.000 immature neurons were plated directly on top of the electrode arrays and cells were allowed to settle for 20min after which cells were cultured in 1ml N2B27 medium for up to 3 month. Medium was changed every 2 days. Dependent on the intensity of detected field potentials arrays were measured 2-4 times a week. Data were recorded for 300 sec. per session. Data were acquired using AxIS software and analyzed using NeuroMetrix and Prism GraphPad.

## Author contribution

RR conceived and performed all experiments, analyzed the data and wrote the manuscript. JG helped with reprogramming of iPSC lines. CS designed CRISPR plasmids for isogenic 22qDS lines. RJ conceived the experiments and wrote the manuscript.

## Acknowledgements

We thank R. Alagappan, D. Fu and T. Lungjangwa for their technical support with human PSC cultures. We would like to thank P. Wisniewski, C. Zollo, C. Arano and M. Ly of the Whitehead Institute FACS-core facility. J. Love and S. Gupta of the Whitehead Genome Technology Core helped with RNA sequencing. We would like to thank all members of the Jaenisch laboratory for helpful discussions and comments on the manuscript. This work was supported by a grant from the Simons Foundation to the Simons Center for the Social Brain at MIT. R.J. was supported by NIH grants 1R01NS088538-01, HD 045022 and 2R01MH104610-15.

## Supplementary Figure legends

**Figure S1: Patient-specific cortical neurons show synaptic and intracellular signaling phenotype. (related to figure 1)**

(A) Scheme of post-synaptic density (PSD) proteins and down-stream signaling pathways.

(B) Selected PSD protein analysis comparing PSC-derived control and two independent patient neuron cultures.

(C) Selected down-stream pathway protein analysis comparing PSC-derived control and two independent patient neuron cultures.

**Figure S2: PAX6-GFP reporter lines allow specific purification of human PSC-derived neural progenitors. (related to figure 2)**

(A) Sequencing of the 2^nd^ allele of three independent clones.

(B) FACS purification of three independent clones plus negative control after NPC differentiation.

(C) Comparative gene expression profiles: global vs. differentially expressed genes.

(D) Comparative transcript profiling of a subset of NPC markers.

(E) qRT-PCR validation of a subset of NPC markers. Error bars represent SEM.

(F) Gene expression up-stream and across the 22q13 deletion. Data are represented as log2 RPM values.

## References

Aldinger, K.A., Kogan, J., Kimonis, V., Fernandez, B., Horn, D., Klopocki, E., Chung, B., Toutain, A., Weksberg, R., Millen, K.J., et al. (2013).Cerebellar and posterior fossa malformations in patients with autism-associated chromosome 22q13 terminal deletion. Am. J. Med. Genet. A 161A, 131–136.

Bidinosti, M., Botta, P., Kruttner, S., Proenca, C.C., Stoehr, N., Bernhard, M., Fruh, I., Mueller, M., Bonenfant, D., Voshol, H., et al. (2016).CLK2 inhibition ameliorates autistic features associated with SHANK3 deficiency. Science (80-.).

Bonaglia, M.C., Giorda, R., Beri, S., De Agostini, C., Novara, F., Fichera, M., Grillo, L., Galesi, O., Vetro, A., Ciccone, R., et al. (2011).Molecular mechanisms generating and stabilizing terminal 22q13 deletions in 44 subjects with Phelan/McDermid syndrome. PLoS Genet.7, e1002173.

Bozdagi, O., Tavassoli, T., and Buxbaum, J.D. (2013).Insulin-like growth factor-1 rescues synaptic and motor deficits in a mouse model of autism and developmental delay. Mol. Autism 4, 9.

Calderon de Anda, F., Rosario, A.L., Durak, O., Tran, T., Gräff, J., Meletis, K., Rei, D., Soda, T., Madabhushi, R., Ginty, D.D., et al. (2012).Autism spectrum disorder susceptibility gene TAOK2 affects basal dendrite formation in the neocortex. Nat. Neurosci. 15, 1022–1031.

Chambers, S.M., Fasano, C. a, Papapetrou, E.P., Tomishima, M., Sadelain, M., and Studer, L. (2009).Highly efficient neural conversion of human ES and iPS cells by dual inhibition of SMAD signaling. Nat. Biotechnol. 27, 275–280

Coffey, E.T. (2014a). Nuclear and cytosolic JNK signalling in neurons. Nat. Rev. Neurosci. 15, 285–299.

Coffey, E.T. (2014b). Nuclear and cytosolic JNK signalling in neurons. Nat. Rev. Neurosci. 15, 285–299.

Cong, L., Ran, F.A., Cox, D., Lin, S., Barretto, R., Habib, N., Hsu, P.D., Wu, X., Jiang, W., Marraffini, L.A., et al. (2013).Multiplex genome engineering using CRISPR/Cas systems. Science 339, 819–823.

Durand, C.M., Betancur, C., Boeckers, T.M., Bockmann, J., Chaste, P., Fauchereau, F., Nygren, G., Rastam, M., Gillberg, I.C., Anckarsäter, H., et al. (2007).Mutations in the gene encoding the synaptic scaffolding protein SHANK3 are associated with autism spectrum disorders. Nat. Genet. 39, 25–27.

Georgala, P.A., Carr, C.B., and Price, D.J. (2011).The role of Pax6 in forebrain development. Dev. Neurobiol. 71, 690–709.

Giza, J., Urbanski, M.J., Prestori, F., Bandyopadhyay, B., Yam, A., Friedrich, V., Kelley, K., D’Angelo, E., and Goldfarb, M. (2010).Behavioral and cerebellar transmission deficits in mice lacking the autism-linked gene islet brain-2. J. Neurosci. 30, 14805–14816.

Hockemeyer, D., Soldner, F., Beard, C., Gao, Q., Mitalipova, M., DeKelver, R.C., Katibah, G.E., Amora, R., Boydston, E.A., Zeitler, B., et al. (2009).Efficient targeting of expressed and silent genes in human ESCs and iPSCs using zinc-finger nucleases. Nat. Biotechnol. 27, 851–857.

Jiang, Y.-H., and Ehlers, M.D. (2013).Modeling autism by SHANK gene mutations in mice. Neuron 78, 8–27.

Kolevzon, A., Bush, L., Wang, A.T., Halpern, D., Frank, Y., Grodberg, D., Rapaport, R., Tavassoli, T., Chaplin, W., Soorya, L., et al. (2014).A pilot controlled trial of insulin-like growth factor-1 in children with Phelan-McDermid syndrome. Mol. Autism 5, 54.

Koushika, S.P. (2008).“JIP”ing along the axon: the complex roles of JIPs in axonal transport. Bioessays 30, 10–14.

Kukekov, N. V, Xu, Z., and Greene, L.A. (2006).Direct interaction of the molecular scaffolds POSH and JIP is required for apoptotic activation of JNKs. J. Biol. Chem. 281, 15517–15524.

Lengner, C.J., Gimelbrant, A.A., Erwin, J.A., Cheng, A.W., Guenther, M.G., Welstead, G.G., Alagappan, R., Frampton, G.M., Xu, P., Muffat, J., et al. (2010).Derivation of pre-X inactivation human embryonic stem cells under physiological oxygen concentrations. Cell 141, 872–883.

Li, Y., Wang, H., Muffat, J., Cheng, A.W., Orlando, D.A., Lovén, J., Kwok, S.-M., Feldman, D.A., Bateup, H.S., Gao, Q., et al. (2013). Global transcriptional and translational repression in human-embryonic-stem-cell-derived rett syndrome neurons. Cell Stem Cell 13, 446–458.

Lu, C., Chen, Q., Zhou, T., Bozic, D., Fu, Z., Pan, J.Q., and Feng, G. (2016).Micro-electrode array recordings reveal reductions in both excitation and inhibition in cultured cortical neuron networks lacking Shank3. Mol. Psychiatry 21, 159–168.

Maden, C.H., Gomes, J., Schwarz, Q., Davidson, K., Tinker, A., and Ruhrberg, C. (2012).NRP1 and NRP2 cooperate to regulate gangliogenesis, axon guidance and target innervation in the sympathetic nervous system. Dev. Biol. 369, 277–285.

Mali, P., Yang, L., Esvelt, K.M., Aach, J., Guell, M., DiCarlo, J.E., Norville, J.E., and Church, G.M. (2013).RNA-guided human genome engineering via Cas9. Science 339, 823–826.

Mei, Y., Monteiro, P., Zhou, Y., Kim, J.-A., Gao, X., Fu, Z., and Feng, G. (2016).Adult restoration of Shank3 expression rescues selective autistic-like phenotypes. Nature.

Mooney, L.M., and Whitmarsh, A.J. (2004).Docking interactions in the c-Jun N-terminal kinase pathway. J. Biol. Chem. 279, 11843–11852.

Osumi, N., Shinohara, H., Numayama-Tsuruta, K., and Maekawa, M. (2008).Concise review: Pax6 transcription factor contributes to both embryonic and adult neurogenesis as a multifunctional regulator. Stem Cells 26, 1663–1672.

Pasterkamp, R.J. (2012).Getting neural circuits into shape with semaphorins. Nat. Rev. Neurosci. 13, 605–618.

Peça, J., Feliciano, C., Ting, J.T., Wang, W., Wells, M.F., Venkatraman, T.N., Lascola, C.D., Fu, Z., and Feng, G. (2011).Shank3 mutant mice display autistic-like behaviours and striatal dysfunction. Nature 472, 437–442.

Phelan, M.C. (2008).Deletion 22q13.3 syndrome. Orphanet J. Rare Dis. 3, 14.

Sarasua, S.M., Boccuto, L., Sharp, J.L., Dwivedi, A., Chen, C.-F., Rollins, J.D., Rogers, R.C., Phelan, K., and DuPont, B.R. (2014).Clinical and genomic evaluation of 201 patients with Phelan-McDermid syndrome. Hum. Genet. 133, 847–859.

Shcheglovitov, A., Shcheglovitova, O., Yazawa, M., Portmann, T., Shu, R., Sebastiano, V., Krawisz, A., Froehlich, W., Bernstein, J. a, Hallmayer, J.F., et al. (2013).SHANK3 and IGF1 restore synaptic deficits in neurons from 22q13 deletion syndrome patients. Nature 503, 267–271.

Shi, Y., Kirwan, P., and Livesey, F.J. (2012a). Directed differentiation of human pluripotent stem cells to cerebral cortex neurons and neural networks. Nat. Protoc. 7, 1836–1846.

Shi, Y., Kirwan, P., Smith, J., Robinson, H.P.C., and Livesey, F.J. (2012b).Human cerebral cortex development from pluripotent stem cells to functional excitatory synapses. Nat. Neurosci. 15, 477–486, S1.

Soldner, F., and Jaenisch, R. (2012).Medicine. iPSC disease modeling. Science 338, 1155–1156.

Soldner, F., Laganière, J., Cheng, A.W., Hockemeyer, D., Gao, Q., Alagappan, R., Khurana, V., Golbe, L.I., Myers, R.H., Lindquist, S., et al. (2011).Generation of Isogenic Pluripotent Stem Cells Differing Exclusively at Two Early Onset Parkinson Point Mutations. Cell 146, 318–331

Soldner, F., Stelzer, Y., Shivalila, C.S., Abraham, B.J., Latourelle, J.C., Barrasa, M.I., Goldmann, J., Myers, R.H., Young, R.A., and Jaenisch, R. (2016).Parkinson-associated risk variant in distal enhancer of α-synuclein modulates target gene expression. Nature 533, 95–99.

Takahashi, K., and Yamanaka, S. (2013).Induced pluripotent stem cells in medicine and biology. Development 140, 2457–2461.

Tiwari, V.K., Stadler, M.B., Wirbelauer, C., Paro, R., Schübeler, D., and Beisel, C. (2011).A chromatin-modifying function of JNK during stem cell differentiation. Nat. Genet. 44, 94–100.

Tran, T.S., Rubio, M.E., Clem, R.L., Johnson, D., Case, L., Tessier-Lavigne, M., Huganir, R.L., Ginty, D.D., and Kolodkin, A.L. (2009).Secreted semaphorins control spine distribution and morphogenesis in the postnatal CNS. Nature 462, 1065–1069.

Uchino, S., and Waga, C. (2013).SHANK3 as an autism spectrum disorder-associated gene. Brain Dev. 35, 106–110.

Warren, L., Manos, P.D., Ahfeldt, T., Loh, Y.-H., Li, H., Lau, F., Ebina, W., Mandal, P.K., Smith, Z.D., Meissner, A., et al. (2010).Highly efficient reprogramming to pluripotency and directed differentiation of human cells with synthetic modified mRNA. Cell Stem Cell 7, 618–630.

Whitmarsh, A.J. (2006).The JIP family of MAPK scaffold proteins. Biochem. Soc. Trans. 34, 828–832.

Yi, F., Danko, T., Botelho, S.C., Patzke, C., Pak, C., Wernig, M., and Südhof, T.C. (2016).Autism-associated SHANK3 haploinsufficiency causes Ih channelopathy in human neurons. Science 352, aaf2669

